# A reporter mouse model to map RIG-I expression and function *in vivo*

**DOI:** 10.64898/2026.11.04.764288

**Authors:** Stephen A. Ehrenzeller, Amy C. Hinesley, Sergio Borunda, Suhan Liu, Daniel O. Richards, Dong-il Kwon, Timothy M. Mehrmann, William M. Phillbrick, Daniel T. Thoresen, Richard A. Flavell, Anna Marie Pyle, Akiko Iwasaki

## Abstract

The cytosolic RNA sensor RIG-I is a critical component of the innate immune response to RNA viruses, yet few tools exist to track RIG-I *in vivo*. Herein, we report the development of a RIG-I reporter mouse, termed RIG-I^mScar^, that enables fluorescence imaging and tractable immunoprecipitation of RIG-I *in vivo*. These mice revealed enrichment of RIG-I expression in circulatory, lymphoid, and mucosal tissues and broad expression across immune cell types. RIG-I itself is upregulated in response to IFN. To distinguish RIG-I sentinel cells, which are programmed to express RIG-I at steady state, from IFN-responder cells, we crossed RIG-I^mScar^ mice with Mx1^GFP^ mice. Blocking IFNAR with a monoclonal antibody showed that basal RIG-I expression in immune cells was largely independent of type I IFN signaling, although Ly6C^hi^ monocytes required IFNAR for RIG-I expression. To probe functional responses *in vivo*, we intranasally administered a RIG-I-specific agonist, SLR14. Mapping RIG-I expression across the lung identified alveolar macrophages as a critical RIG-I sensor cell population, and depletion of alveolar macrophages, but not neutrophils, slightly impaired lung IFN responses. Together, these findings support the use of the RIG-I^mScar^ model to probe RIG-I dynamics *in vivo* and predict functional sensor cell populations across tissues.

## INTRODUCTION

The innate immune system relies on a catalog of pattern-recognition receptors (PRRs) to detect conserved molecular motifs of potential pathogens and initiate an immune response.^1^ Among these, retinoic acid-inducible gene I (RIG-I) is a cytosolic RNA sensor involved in the recognition and control of RNA viruses. RIG-I selectively binds blunt-ended double-stranded RNA (dsRNA) species bearing terminal 5’-tri- or diphosphate groups, which are common features of viral genomes and their replication intermediates.^2–6^ Upon RNA recognition, RIG-I signals through mitochondrial antiviral signaling protein (MAVS) to activate key transcriptional regulators of the innate immune response such as IRF3 and NF-KB, inducing the expression of interferon stimulated genes (ISGs; e.g., *Mx1*), pro-inflammatory cytokines, and type I interferons (IFNs).^6^ RIG-I itself is also an ISG, although interferon-induced RIG-I species are not involved in canonical MAVS-dependent antiviral signaling.^7^ The essential and non-redundant role of RIG-I in host antiviral defense has been well-established, as RIG-I-deficient mice exhibit enhanced susceptibility to RNA viruses such as Japanese encephalitis virus (JEV)^8^ and West Nile virus (WNV).^9^ Moreover, pharmacologic targeting of RIG-I via synthetic oligonucleotides such as stem-loop RNAs (e.g., SLR14) has shown pre-clinical promise as a prophylactic and therapeutic antiviral^10–12^ and intratumoral immunotherapy,^13–15^ underscoring RIG-I as a potential target for innate immune modulation in infectious diseases and cancer.

Despite these advances, cell- and tissue-specific patterns of RIG-I expression and function *in vivo* remain incompletely understood. RIG-I dynamics are classically modeled in *in vitro* systems, which are adequate tools to investigate structure-function relationships and subcellular organization in RIG-I signaling. However, these systems often rely on select immortalized cell lines and artificial expression platforms that fail to capture the spatial and cellular heterogeneity of RIG-I expression and function within tissues. Indeed, although RIG-I is expressed across a broad range of cell types and tissues,^16,17^ the regulatory mechanisms governing its expression, the contributions of distinct cell populations to antiviral immunosurveillance, and the extent to which RIG-I signaling mechanisms are conserved across cell types remain poorly understood. Defining these patterns will help elucidate how productive antiviral immune responses are orchestrated across tissues and requires an approach that enables direct visualization and characterization of endogenous RIG-I expression across tissues and cell populations.

In the present work, we report the generation of a RIG-I reporter mouse model, RIG-I^mScar^, which expresses a mScarlet-3xHA-RIG-I fusion construct at the endogenous RIG-I (Ddx58) locus, enabling robust monitoring of physiological RIG-I expression *in vivo*. Using this model, we demonstrate enriched RIG-I expression at lymphoid and mucosal tissues, with RIG-I broadly expressed across immune cell subsets. To distinguish homeostatic sentinel cells that detect cytosolic RNA via constitutive RIG-I expression from IFN-responder cells, we crossed RIG-I^mScar^ mice with Mx1^GFP^ mice, enabling us to distinguish cells with steady-state RIG-I expression (mScarlet^+^) from those induced by type I IFN (GFP^+^). Using both reporter lines, we examined RIG-I expression profiles at steady state and candidate sensor cells after agonist administration *in vivo*.

## RESULTS

### Development and Characterization of the RIG-I^mScar^ Model

To generate mice with RIG-I bearing a mScarlet fluorescent reporter and hemagglutinin (HA) pull-down tag (hereafter referred to as “RIG-I^mScar^ mice”), we inserted mScarlet, HA, and GGS linker sequences at the 5’ terminus of the first exon of the murine RIG-I gene via CRISPR-mediated, homology-directed integration.^18^ The resulting RIG-I construct is expressed under the control of the endogenous regulatory elements of the RIG-I gene (Ddx58) and features a mScarlet-HA fusion linked to the N-terminus of the native RIG-I protein via a 6x GGS linker (Fig. 1A). Successful insertion without introduction of off-target mutations was confirmed via whole-genome sequencing (Fig. 1B).

**Figure 1.**
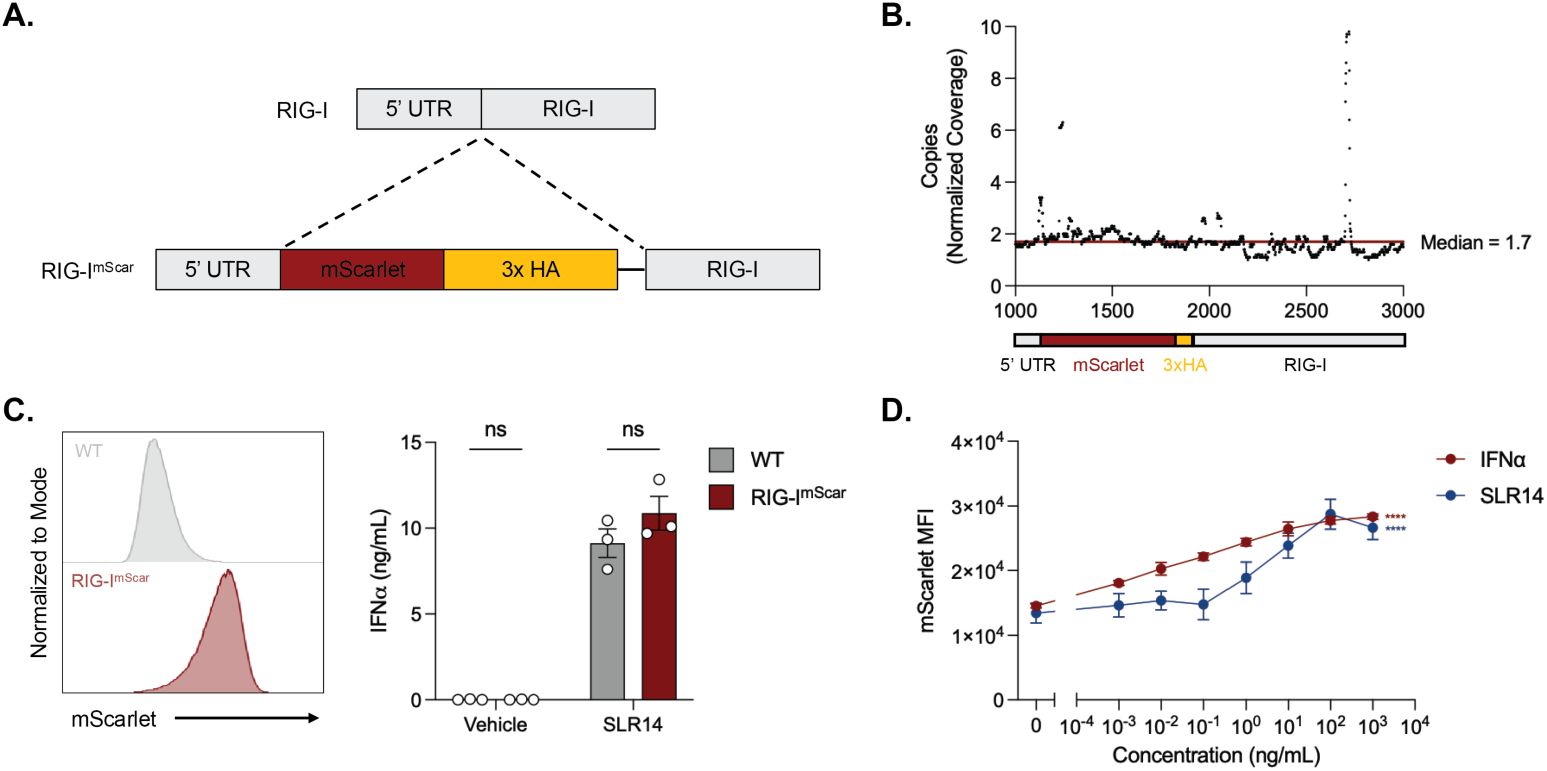
Validation of RIG-I^mScar^ construct expression and function. (A) Schematic of RIG-I^mScar^ construct, featuring the mScarlet and 3x HA tag insertions downstream of the endogenous 5’ UTR of the RIG-I gene. The mScarlet and HA tags are linked to the native RIG-I protein via 6x GGS linker. (B) Validation of mScarlet-3xHA cassette insertion via whole genome sequencing (WGS). Genomic DNA was isolated from spleen lysates of RIG-I mScar mice and subject to Illumina WGS. Reads were aligned to the expected mScarlet-3xHA-RIG-I locus, and copy number was calculated as calculated by normalizing the number of aligned reads to the average coverage of the entire sample. (C) Representative flow cytometry histogram of mScarlet signal in bone-marrow derived macrophages (BMDMs) cultured from C57BL/6 and RIG-I^mScar/mScar^ mice (left). Supernatant IFNα concentrations after 24-hour culture with SLR14 complexed with lipofectamine (right). Data were analyzed via one-way ANOVA with Šidák’s multiple comparison test. (D) mScarlet MFI in BMDMs cultured with the indicated concentrations of murine IFNα or SLR14 complexed with lipofectamine. Data were analyzed via ordinary two-way ANOVA with Dunnett’s multiple comparison test. All data are presented as mean ± SEM (n = 3/ea.).

The N-terminus of RIG-I features two caspase recruitment domains (CARDs) that facilitate the interaction of RIG-I with its signaling partner MAVS,^6^ and it is possible that insertion of the mScarlet-HA tag disrupts RIG-I-MAVS interactions and downstream signal transduction. To determine potential functional consequences of the mScarlet-3xHA-RIG-I construct in primary cells, we generated bone marrow-derived macrophages (BMDMs) from C57BL/6 (WT) or RIG-I^mScar^ mice. Basal expression of the mScarlet tag was readily distinguishable in RIG-I^mScar^ BMDMs (Fig. 1C). BMDMs treated with the selective RIG-I ligand SLR14 showed comparable IFN-α secretion between WT and RIG-I^mScar^ BMDMs, indicating preserved function of the mScarlet-3xHA-RIG-I construct (Fig. 1D). mScarlet median fluorescence intensity (MFI) was also enhanced in a dose-dependent manner following SLR14 treatment (Fig. 1D). These effects were mirrored with type I IFN (IFN-α) treatment, in keeping with the known role of RIG-I as an ISG downstream of IFNAR.^19^ Thus, the RIG-I^mScar^ remained functional within BMDM *ex vivo*.

### RIG-I-mScarlet Expression is Variable Across Organs and Cell Types

We next sought to profile RIG-I-mScarlet expression *in vivo*. Analysis of major tissues revealed distinct expression patterns, with RIG-I-mScarlet MFI primarily enriched in circulation (blood), secondary lymphoid (spleen, lymph node) and mucosal (lung, small intestine lamina propria (SILP), female reproductive tract (FRT)) tissues, with low MFI detected in the brain, kidney and the bone marrow (Fig. 2A). Given that lymphoid tissues feature dense immune cell networks and that mucosal surfaces are the primary portal of entry for pathogens,^20^ these findings suggest that RIG-I expression may be enriched in tissues with active immunosurveillance and frontline host defense. RIG-I-mScarlet MFI was remarkably low in the brain and kidney, likely reflecting the comparably low frequency of mature immune cells in these tissues. RIG-I expression was similarly dim in the bone marrow, perhaps reflecting weaker expression in stromal or hematopoietic cell types. Analysis of major immune cell types demonstrated distinct expression patterns as well, although these were largely conserved across tissues (Fig. 2B). In myeloid lineages, we observed the greatest frequency of RIG-I-mScarlet^+^ cells in macrophages, neutrophils, and Ly6C^lo^ monocytes. Surprisingly, lymphoid lineages exhibited comparably higher expression, with enrichment of RIG-I-mScarlet^+^ cells in CD4^+^ and CD8^+^ T cells. Because these cells are not thought to participate in viral sensing and innate immune defense, RIG-I may serve non-classical functions in these cell types. To validate these findings, we compared RIG-I-mScarlet expression to pre-analyzed, published single-cell RNA-sequencing data from the Immunological Genome Project (ImmGen) for RIG-I expression.^21^ In age- and sex-matched WT C57BL/6 mice, RIG-I transcript expression was enriched across splenic immune cells (Fig. S2A), and RIG-I-mScarlet signal correlated positively (R^2^ = 0.3181) and significantly (p = 0.0002) with transcript level among the same cell types (Fig. S2B). Thus, the RIG-I^mScar^ model faithfully reflects endogenous RIG-I expression.

**Figure 2.**
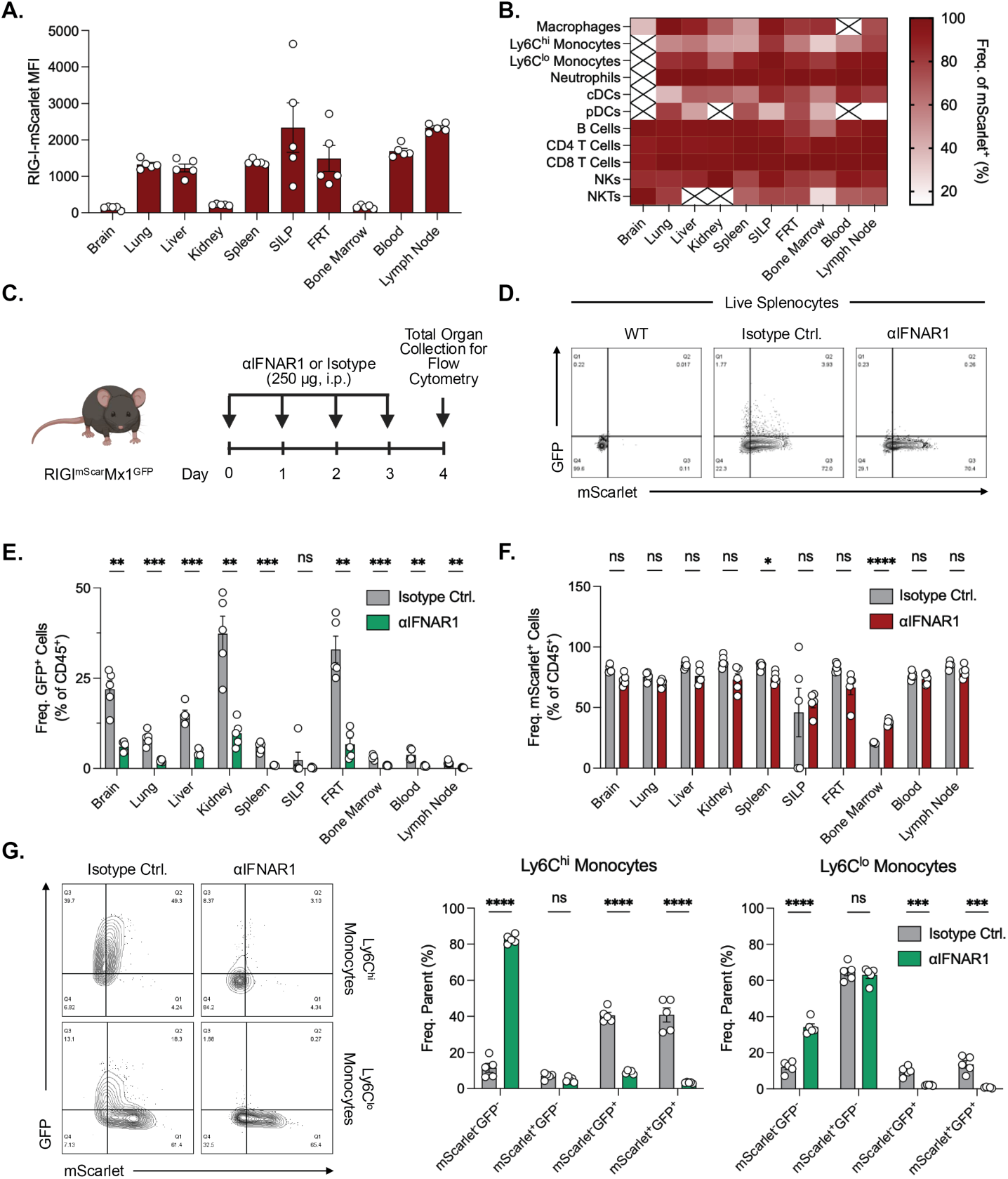
Whole organism mapping of RIG-I^mScar^ expression *in vivo*. (A) mScarlet MFI of total live cells isolated from the indicated organs of RIG-I^mScar/mScar^ mice. (B) Heat map displaying the frequency of mScarlet^+^ cells in the indicated myeloid and lymphoid immune cell populations across various organs. (C) Experimental design. RIG-I^mScar/+^Mx1^GFP/+^ were administered 250 μg of isotype control or αIFNAR1 monoclonal antibody intraperitoneally for 3 days prior to organ harvest. Immune cell GFP and mScarlet MFI were assessed via flow cytometry. (D) Representative flow cytometry plots of total live splenocytes from WT C57BL/6J (left) or RIG-I^mScar/+^Mx1^GFP/+^ mice treated with isotype control (middle) or αIFNAR1 antibodies (right). (E) Frequency of GFP^+^ CD45^+^ cells in the indicated organs following treatment with isotype control or αIFNAR1 antibodies. (F) Frequency of mScarlet^+^ CD45^+^ cells in the indicated organs following treatment with isotype control or αIFNAR1 antibodies. (G) Representative flow cytometry plots of Mx1-GFP and RIG-I-mScarlet signal in Ly6C^hi^ and Ly6C^lo^ monocytes following αIFNAR1 blockade (left) and frequencies of GFP^+^ and mScarlet^+^ cells (right). GFP^+^ and mScarlet^+^ populations were determined via gating against representative WT controls. All data were analyzed via multiple unpaired t-tests with Holm-Šidák’s multiple comparisons test. All data are presented as mean ± SEM (n = 4-5/ea.). SILP = small intestine lamina propria; FRT = Female reproductive tract. cDCs = conventional dendritic cells; pDCs = plasmacytoid dendritic cells.

### Homeostatic RIG-I-mScarlet Expression is Largely Independent of Type I IFN Signaling

Despite the established role of RIG-I in mediating protective antiviral immune responses, its expression dynamics *in vivo* remain incompletely understood. Proteomic^16,17^ and transcriptomic^21^ analyses, as well as our present findings, corroborate RIG-I expression across most tissues and nucleated cell types, but how such expression patterns are established and regulated remains unclear. Defining the mechanisms that establish constitutive RIG-I expression may therefore clarify how cells are programmed for antiviral immune surveillance independent of acute IFN stimulation. As such, we generated a RIG-I^mScar^Mx1^GFP^ dual-reporter system for simultaneous tracking of RIG-I expression and Mx1-GFP to report on ISG induction.^22^

At steady state, Mx1-GFP was expressed in CD45^+^ cell populations in most organs tested, with particularly high expression found in the brain, kidney, and FRT (Fig. 2E), while RIG-I-mScarlet signals were observed at similar levels across all organs except for the bone marrow, which had lower expression (Fig. 2F). We treated these mice with either interferon alpha/beta receptor blockade (αIFNAR1) or isotype control antibody to assess IFN-dependent constitutive RIG-I expression (Fig. 2C). Compared with RIG-I-mScarlet, basal Mx1-GFP expression was low, and this expression was largely abrogated following αIFNAR1 treatment (Fig. 2D–E). The frequency of Mx1-GFP^+^ immune cells was significantly reduced by IFNAR blockade of most organs analyzed, with the strongest reduction observed in the brain, kidney, and FRT (Fig. 2E). In contrast, the frequency of RIG-I-mScarlet^+^ immune cells remained stable across tissues (Fig. 2F), and these findings were largely conserved across immune cell types (Fig. S3), suggesting RIG-I expression is largely maintained in an IFN-independent manner. Nevertheless, cell-specific distinctions emerged within the myeloid compartment. Consistent with previous reports,^22^ Ly6C^hi^ monocytes (i.e., “classical” or “inflammatory”) exhibited high basal Mx1-GFP expression, whereas Ly6C^lo^ monocytes (i.e., “non-classical” or “patrolling”) exhibited low basal Mx1-GFP expression. In keeping, RIG-I-mScarlet was co-expressed with Mx1-GFP in Ly6C^hi^ monocytes, whereas RIG-I expression was mostly decoupled from Mx1-GFP expression in Ly6C^lo^ monocytes (Fig. 2G, grey). Following αIFNAR1 treatment, RIG-I-mScarlet expression in Ly6C^hi^ monocytes was largely lost (>90% reduction), whereas expression in Ly6C^lo^ monocytes remained stable in a Mx1-GFP-negative population. These findings indicate distinct modes of RIG-I regulation in monocytes, where expression in Ly6C^hi^ monocytes is likely maintained through a constitutive ISG program, while expression in Ly6C^lo^ monocytes is largely maintained by an IFN-independent transcriptional program.

### RIG-I^mScar^ Model Identifies Key Role of Airway Immune Cells in RNA Sensing

*In vivo* mapping of RIG-I expression might also help identify functional RIG-I sensor cells. For example, the respiratory mucosa must be equipped to sense and respond rapidly to highly transmissible RNA viruses such as influenza viruses, coronaviruses, and respiratory syncytial virus (RSV). Our present findings indicate comparatively high RIG-I expression in the lung, and studies of RIG-I-deficient mice demonstrate increased susceptibility and impaired control of respiratory infection in the absence of RIG-I.^23^ Moreover, our groups previously demonstrated that intravenous delivery of SLR14 in complex with JET-PEI enhanced IFN expression in the lung and conferred therapeutic and prophylactic protection against SARS-CoV-2.^11^ *In vitro* studies have suggested that epithelial cells,^24,25^ macrophages,^26,27^ and dendritic cells^28,29^ are competent RIG-I sensor cells, although their functional relevance *in vivo* remains unclear. As such, we used the RIG-I^mScar^ mouse to probe RIG-I expression in the lung and predict functional RIG-I sensor cell populations.

We first assessed basal RIG-I-mScarlet expression across candidate immune and stromal cell populations in the lung via flow cytometry. mScarlet MFI was highest in epithelial cells, alveolar macrophages, and neutrophils, consistent with previous *in vitro* studies (Fig. 3A).^24–27,30^ To assess the role of alveolar macrophages in RIG-I-mediated sensing in the lung, we depleted alveolar macrophages with clodronate liposomes prior to intranasal challenge with SLR14 (Fig. 3B). Clodronate liposome treatment resulted in >90% depletion of alveolar macrophages (Fig. S5A). Four hours after challenge, lungs and bronchioalveolar lavage fluid (BALF) were collected to assess local IFN responses. In the lung, other than *Isg15,* expression of a panel of ISGs (*Mx1, Cxcl10, Irf7, Oas1, Ifit1*) showed no significant reduction following alveolar macrophage depletion (Fig. 3E), whereas BALF IFNα protein levels were significantly decreased (Fig. 3F). These deficits were not observed following successful depletion of neutrophils with αLy6G antibodies (Fig. 3G–I, Fig. S5B) despite high RIG-I expression in these cells, suggesting a specific role for alveolar macrophages in RIG-I sensing at the respiratory mucosa. Indeed, these findings are consistent with previous reports of alveolar macrophages as the primary source of lung IFNs in murine RNA virus infections.^31,32^ Importantly, ISG responses remained elevated compared to the vehicle control, even with alveolar macrophage depletion, suggesting redundant and/or compensatory sensing by other cell populations such as epithelial cells or dendritic cells.

**Figure 3.**
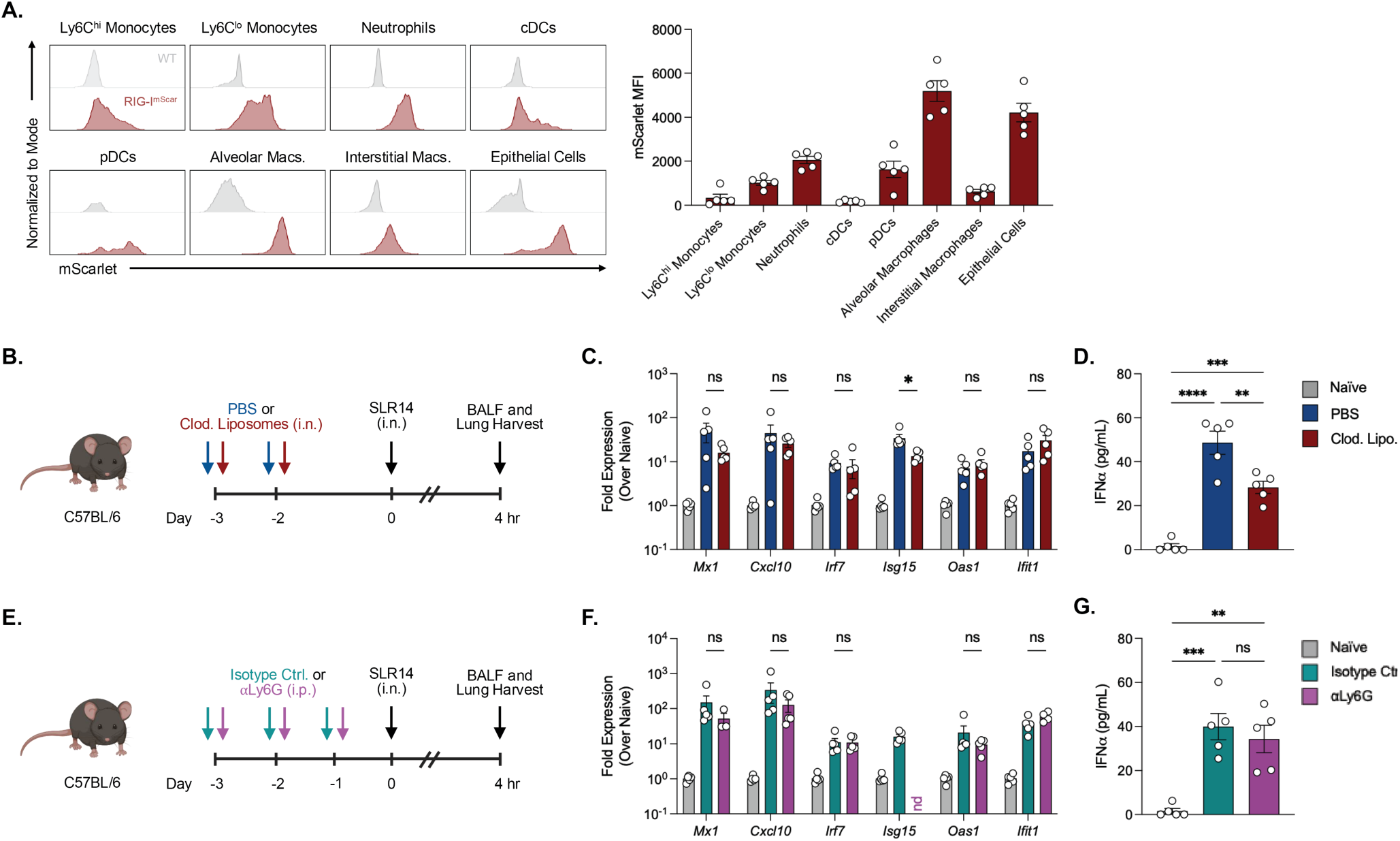
Alveolar macrophages, but not neutrophils, direct lung IFN responses to intranasal SLR14. (A) Representative flow cytometry histograms of mScarlet signal in the indicated lung cell types of WT or RIG-I^mScar/mScar^ mice (left). Corresponding MFIs of the indicated lung cell types (right). (B) Experimental design. C57BL/6J mice were administered PBS or clodronate liposomes intranasally to deplete alveolar macrophages. Mice were then challenged intranasally with SLR14, and BAL and lung tissue were collected 4 hours post-challenge to assess ISG activation and IFN secretion. (C) ISG expression in lung lysates following intranasal SLR14 treatment. (D) IFNα content of BALF following intranasal SLR14 treatment. (E) Experimental design. C57BL/6J mice were administered isotype control or αLy6G antibodies to deplete neutrophils. Mice were then challenged intranasally with SLR14, and BAL and lung tissue were collected 4 hours post-challenge to assess ISG activation and IFN secretion. (F) ISG expression in lung lysates following intranasal SLR14 treatment. (G) IFNα content of BALF following intranasal SLR14 treatment. Expression data were analyzed via multiple unpaired t-tests with Holm-Šidák’s multiple comparisons test, and IFNα data were analyzed via ordinary two-way ANOVA with Šidák’s multiple comparisons test. All data are presented as mean ± SEM (n = 5/ea.). cDCs = conventional dendritic cells; pDCs = plasmacytoid dendritic cells.

## DISCUSSION

The ability to define where innate immune sensors are expressed *in vivo* is fundamental to understanding how tissues establish and maintain antiviral immune surveillance. Although RIG-I is a well-characterized cytosolic RNA sensor, the tissue distribution, cellular regulation, and functional contributions of RIG-I-expressing cells under homeostatic conditions remain incompletely understood.^8,33,34^ In this study, we describe RIG-I^mScar^ mice, a fluorescent reporter model that enables visualization and analysis of RIG-I expression *in vivo*, allowing us to identify candidate sensor populations in their physiological tissue context.

The RIG-I^mScar^ mouse features a mScarlet-3xHA fusion protein knocked in at the 5’-end of the first exon of the RIG-I coding sequence, resulting in the expression of a mScarlet-3xHA-RIG-I construct at the endogenous RIG-I locus. We demonstrated that this reporter does not impair response to the RIG-I-specific ligand SLR14, suggesting retained RIG-I function, and we found that reporter expression is inducible by canonical stimuli such as type I IFN and RIG-I-specific ligands.

RIG-I-mScarlet expression varied across tissues and cell types *in vivo*. It was particularly enriched in lymphoid and mucosal tissues, sites of pathogen exposure, immune activation, and coordination of local and systemic immune responses. Enrichment of RIG-I in these tissues may reflect the importance of maintaining RNA-sensing capacity at sites where viral exposure or dissemination is likely to occur. In mucosal tissues, such enrichment may enable rapid recognition of viral RNA before an infection can become established, whereas in circulating and secondary lymphoid compartments, it may support infection detection and induction of adaptive antiviral immune responses. Within the immune cell compartment, RIG-I expression was enriched in populations with established roles in immune surveillance, including macrophages, neutrophils, and Ly6C^lo^ monocytes.^35^ Unexpectedly, CD4^+^ and CD8^+^ T cells were also found to have relatively high RIG-I expression, suggesting potential functions beyond conventional innate immune sensing. Indeed, emerging evidence suggests RIG-I and RIG-I-like receptors (RLRs) might modulate T cell survival, fitness, and effector function,^36,37^ although whether these processes proceed through canonical sensing of dsRNA remains unclear. Additionally, elevated reporter expression alone does not necessarily indicate that tissues and cell populations have higher RIG-I-dependent sensing capacity, which also depends on ligand accessibility, cell activation state, and downstream signaling.

RIG-I expression was largely preserved across cell types and tissues following an IFNAR1 blockade despite effective suppression of the Mx1-GFP reporter *in vivo*, suggesting IFNAR-independent control of RIG-I expression. However, cell-specific differences were revealed in monocytes, where RIG-I expression was drastically reduced in Ly6C^hi^ monocytes and only moderately reduced in Ly6C^lo^ monocytes. These findings are in keeping with previous reports of IFN-dependent differentiation of Ly6C^hi^ monocytes^38^ and suggest an alternative transcription program regulating RIG-I expression in these cells. Future studies will seek to assess the transcriptional and epigenetic mechanisms by which constitutive IFN stimulus regulates RIG-I expression in these cell types. Moreover, considering most immune cells expressed RIG-I in an IFN-independent fashion, it will be equally relevant to understand how transcriptional programs poise these cells for immunosurveillance and how these programs might adapt to acute and persistent RIG-I stimulation. Additionally, the source of IFN-I responsible for driving the Mx1-GFP expression found at steady state in our SPF mouse condition remains unidentified.

Because the lung is a common site of RNA virus infection, we sought to use the RIG-I^mScar^ model to investigate functional RIG-I sensor cells in an SLR14 challenge model. Baseline RIG-I-mScarlet expression was strongest in epithelial cells, alveolar macrophages, and plasmacytoid dendritic cells, consistent with their established roles as sensors of viral infection.^24–29^ Surprisingly, RIG-I expression was also high in neutrophils, whose role in antiviral immunity is less well understood.^39^ Previous reports demonstrated that RIG-I is partially sequestered into secretory vesicles in human neutrophils,^40^ suggesting a potential non-canonical role of RIG-I. Despite these findings, depletion of neutrophils with αLy6G antibodies had no effect on lung IFN responses to intranasal SLR14 challenge. Rather, depletion of alveolar macrophages with clodronate liposomes resulted in reduced IFNα secretion in BALF. These findings are consistent with previous reports of alveolar macrophages as key antiviral sentinels and interferon-secreting cells in the lung.^31,32^ Nevertheless, the effects of depletion were only partial, suggesting redundancies with alternative cell types. Indeed, our previous reports demonstrated that epithelial cells were the primary pulmonary target of intravenous SLR14 administration,^11^ and our immunofluorescence data corroborate high expression of RIG-I in these cells.

Collectively, these findings highlight the potential of the RIG-I^mScar^ model to monitor RIG-I biology in physiological contexts. The mScarlet reporter offers a tractable system to visualize RIG-I across diverse cell types, which may aid discovery of RIG-I sensor cells *in vivo* or reveal novel cell-specific RIG-I signaling dynamics in primary cells *ex vivo*. Additionally, the 3xHA tag enables the pull-down of RIG-I complexes from tissue, providing an opportunity to identify novel RNA ligands and RIG-I binding partners *in vivo*. Although the present work establishes the model and points to candidate sensor populations, future applications of this model will enable a better understanding of RIG-I biology in infection, autoimmune disease, cancer, and aging.

## MATERIALS AND METHODS

### Mice

Mouse studies were conducted according to National Institutes of Health (NIH) guidelines and the Yale Institutional Animal Care and Use Committee. C57BL/6J (Strain #000664) and B6.Cg-Mx1tm1.1Agsa/J (Mx1GFP, Strain #033219) mice were purchased from Jackson Laboratory and bred in-house. C57BL/6N (Strain #027) mice were purchased from Charles River and bred in-house. Mice were maintained under specific pathogen-free conditions in a temperature and humidity-controlled room with a 12-hour light/12-hour dark cycle and continuous access to food and water. All studies used 6-to 8-week-old mice.

RIG-I^mScar^ mice were generated via CRISPR/Cas9-mediated homology-directed repair (HDR) essentially as previously described,^18^ except for the use of Cas9 nuclease protein (New England Biolabs). The sgRNA and homology arms were designed to target insertion of the mScarlet-3xHA-6xGGS HDR cassette (synthesized as a single-stranded Megamer from Integrated DNA Technologies) between the 5’ UTR and exon 1 of RIG-I. This configuration results in a continuous coding region between the reporter and RIG-I sequences under the control of the endogenous regulatory elements. Reporter insertion and orientation in the founders was confirmed via nested PCR, and whole-genome sequencing was performed on the progeny to verify faithful inheritance and assess the presence of off-target mutations (Fig. 1B). Genotyping was performed with the following primer sets: RIG-I-WT F: 5’-CTGGAGCCACAGAATGACA-3’; RIG-I-WT R: 5’-CGAGCCAGGAACTCATGTAGC-3’; RIG-I-mScarlet F: 5’-GGTCCTTGCGGTTCAGTGGC-3’; RIG-I-mScarlet R: 5’-CTGCCTCGCCCTTGCTCACC-3’.

RIG-I^mScar^ and Mx1^GFP^ mice were crossed to generate RIG-I^mScar^Mx1^GFP^ mice, and the heterozygous F1 progeny were used for all experiments.

### Whole Genome Sequencing

Spleens were harvested from 8-week-old female mScarlet-3xHA-RIG-I mice. DNA was subsequently isolated via phenol-chloroform extraction and submitted to SeqCenter (Pittsburgh, PA, USA) for whole genome sequencing. Illumina sequencing libraries were prepared using the tagmentation-based and PCR-based Illumina DNA Kit and custom IDT 10bp unique dual indices (UDI) with a target insert size of 280 bp. Illumina sequencing was performed on an Illumina NovaSeq X Plus sequencer in one or more multiplexed shared-flow-cell runs to produce 2×151bp paired-end reads with an average coverage of ∼56x. Demultiplexing, quality control and adapter trimming was performed with bcl-convert1 (v4.2.4). The paired-end reads were aligned to the expected mScarlet-3xHA-RIG-I locus, using the SAMtools package (v1.24.0) (https://github.com/samtools/samtools) alignment tools with default options for index and align. Sequencing data have been deposited in the Sequence Read Archive (SRA) database under accession PRJNA1556204.

### Protein Extraction and Immunoprecipitation

Mouse tissue lysates were prepared by homogenizing 0.15–0.3mg of mouse spleen tissue in 5 µL/mg Pierce™ IP Lysis Buffer (ThermoFisher, Cat: 87787). Protein concentration was determined using Pierce™ Bradford Plus Protein Assay Reagent (ThermoFisher, Cat: 23238). 12.5 µL Pierce™ Anti-HA magnetic beads (ThermoFisher, Cat: 88836) were washed with 1mL of PBS-T (PBS + 0.05% Tween 20), then 1 mL of Pierce™ IP Lysis Buffer. 350 µg of protein was added to the washed beads, brought up to 100 µL, and equilibrated overnight at 4°C. Beads were washed according to the manufacturer protocol, then eluted in 40 µL of 1x NuPAGE™ LDS Sample Buffer (ThermoFisher, Cat: NP0008).

Input lysate containing 25 µg of protein, as well as 10 µL of eluate, was electrophoresed under denaturing conditions on 4–12% NuPAGE™ Bis-Tris Mini Protein Gel (Invitrogen, Cat: NP0321BOX) and transferred to PVDF membrane (MilliporeSigma, Cat: IPVH10100). The membranes were blocked for 1 hour with 5% nonfat milk in PBS-T and probed with 1:1000 rabbit anti-mouse RIG-I antibody (LifeTech, Cat: 31460). After washing in PBS-T and probing with horseradish peroxidase-labeled secondary antibody, bound antibodies were identified using SuperSignal™ West Pico PLUS Chemiluminescent Substrate (ThermoFisher, Cat: 34580) and imaged on a BioRad ChemiDoc MP.

### Bone Marrow-Derived Macrophage (BMDM) Differentiation and Culturing

Bone marrow-derived macrophages (BMDMs) were differentiated as previously described.^41^ In brief, femoral and tibial bone marrow was flushed from the marrow cavity and dissociated through a 70 µm filter. The cell suspension was then treated with ammonium-chloride-potassium (ACK) buffer for red blood cell (RBC) lysis prior to culturing. Bone marrow was cultured in DMEM supplemented with 10% FBS), 1% P/S, and 25 ng/mL macrophage colony-stimulating factor (M-CSF) for 7 days. BMDMs were then lifted with 5 mM EDTA and seeded at 5 x 10^4^ cells/well in a 96-well plate for downstream assays.

To compare the response of different RIG-I species to SLR14, BMDMs were differentiated as described above from C57BL/6J and RIG-I^mScar^ bone marrow progenitors. SLR14 was complexed with Lipofectamine 2000 transfection reagent (ThermoFisher, Cat: 11668019) in Opti-PRO serum free media (ThermoFisher, Cat: 12309-050), and BMDMs were cultured with 1 µg/mL of complexed SLR14 for 24 hours. Supernatants were then collected and assessed for IFNα levels via ELISA (Biolegend, Cat: 447904), which was performed following manufacturer’s protocol.

To assess the inducibility of the RIG-I-mScarlet reporter, BMDMs were differentiated as described above from RIG-I^mScar^ bone marrow progenitors. BMDMs were then cultured with a titration of murine IFNα (BioLegend, Cat: 752804), IFNγ (BioLegend, Cat: 575304), or IFNλ (R&D Systems, Cat: 4635-ML-025/CF). BMDMs were cultured with equivalent concentrations of lipofectamine-complexed SLR14 for comparison to a ligand with known RIG-I induction capacity. After 24 hours, cells were washed in PBS, stained with ZombieAqua fixable viability stain diluted in PBS (1:1000) for 10 minutes at room temperature, and fixed in 2% paraformaldehyde (PFA) in PBS for 30 minutes on ice. Flow cytometry data were acquired on a BD FACSymphony A5 Flow Cytometer and analyzed in FlowJo (v10.10.1).

### Mononuclear Cell Isolation & Flow Cytometry

Blood was collected retro-orbitally under anesthesia with isoflurane. Prior to tissue harvest, mice were humanely euthanized under isoflurane and perfused with heparinized PBS. Small intestine lamina propria cell isolation was performed as previously described.^42^ Bone marrow was isolated as described above. Lymph nodes were dissociated in complete RPMI (cRPMI; RPMI 1640 + 10% fetal bovine serum (FBS) + 1% penicillin-streptomycin (P/S)) using a handheld motorized pestle (VWR).

Brains, lungs, livers, kidneys, spleens, and FRTs were harvested, minced with scissors, and incubated in tissue-specific digestion cocktails in cRPMI at 37°C for 45 minutes. Brains were digested with collagenase D (1 mg/mL; Sigma-Aldrich, Cat: 11088882001) and DNase I (30 µg/mL; Sigma-Aldrich, Cat: 10104159001). Lungs were digested with collagenase A (500 µg/mL; Sigma-Aldrich, Cat: 11088793001) and DNase I (50 µg/mL). Livers and kidneys were digested with collagenase IV (200 µg/mL; FisherScientific, 17-104-019) and DNase I (50 µg/mL). Spleens were digested with collagenase D (1 mg/mL) and DNase I (15 µg/mL). FRTs were digested with collagenase D (500 µg/mL), DNase I (30 µg/mL) and dispase II (470 µg/mL; Sigma-Aldrich, Cat: 4942078001). Tissue digestion reactions were quenched with EDTA to 5 mM final concentration and subsequently dissociated through a 70µm filter. Following dissociation, cells were pelleted via centrifugation at 500 x g for 5 minutes at 4°C. Brain pellets were resuspending in 40% Percoll® (Cytiva, Cat: 17-0891-01) in Hank’s Balanced Salt Solution (HBSS) and centrifuged at 580 x g for 15 minutes at room temperature to isolate mononuclear cells. Single-cell suspensions were treated with ACK buffer for RBC lysis. The RBC-free fractions were washed with fluorescence-activated cell sorting (FACS) buffer (PBS + 2% FBS + 1 mM EDTA) prior to downstream assays.

Single-cell suspensions were incubated with either ZombieAqua or ZombieNIR fixable viability dye in PBS (1:1000) for 10 minutes at room temperature. After washing, cells were incubated with primary surface stain including anti-mouse CD16/CD32 Fc block (BioLegend, Cat: 156604) for 1 hour on ice. For the myeloid panel, cells were stained with CD45, CD11b, CD11c, F4/80, I-A/I-E, Ly6-C, Ly-6G, Siglec-F, Siglec-H, and EpCAM. For the lymphoid panel, cells were stained with CD45, CD3ε, CD4, CD8a, CD19, and NK1.1. All antibody stains were prepared at a dilution of 1:200 in FACS buffer. Antibodies details are provided in Table S1. Cells were washed again and fixed with 2% PFA for 30 minutes on ice. Flow cytometry data were acquired on a BD FACSymphony A5 Flow Cytometer and analyzed in FlowJo (v10.10.1). Representative gating strategies are shown in Fig. S1 and Fig. S4.

### *In vivo* Depletions

To block IFNAR1 signaling *in vivo*, mice received 250 µg of anti-mouse IFNAR1 monoclonal antibody (clone MAR1-5A3; BioXCell, Cat: BE0241) intraperitoneally (i.p.) daily for 3 days prior to tissue harvest. Control mice received an isotype matched control IgG1 anti-HRP (Clone HRPN; Bio X Cell, Cat: BE0088). To deplete neutrophils *in vivo*, mice received 150 µg anti-mouse Ly6G (Clone 1A8-CP129; Bio X Cell, CP091) i.p. daily for 3 days prior to SLR14 challenge. Control mice received an isotype matched control IgG2 anti-Chikungunya E1 (Clone CHK-166; Bio X Cell, Cat: BE0496). To deplete alveolar macrophages in the lung, mice were anesthetized with isoflurane and administered 50 µL of clodronate liposome (ApexBio, Cat: K2723) intranasally (i.n.) 72 and 48 hours prior to SLR14 challenge. Control mice received the same volume of PBS, as PBS control liposomes displayed partial depletion of alveolar macrophages.

### SLR14 Challenge

Di-phosphorylated SLR14 was synthesized and purified as previously described.^5,9^ To assess the IFN response to SLR14 in neutrophil-or alveolar macrophage-depleted mice, mice were anesthetized with isoflurane and administered 25 μg of SLR14 in water intranasally. 4 hours later, mice were humanely euthanized. Lungs were collected for RNA extraction and qPCR analysis. Bronchoalveolar lavage fluid (BALF) was collected as previously described^10^ for detection of IFNα levels by ELISA (Biolegend, Cat: 447904), which was performed following manufacturer’s protocol.

### RNA Extraction and Quantification of ISG Expression by qPCR

Lung tissue was placed in a Lysing Matrix D tube (MP Biomedicals, Cat: 6913500) with 1 mL Tri-Zol (Invitrogen, Cat: 155596018) and homogenized for 1 minute at room temperature. RNA was isolated from homogenized mouse tissue samples using Direct-Zol RNA Miniprep kit (Zymo Research, Cat: R2052). cDNA was synthesized from 100 ng of freshly extracted RNA using the iScript cDNA Synthesis kit (Biorad, Cat: 1708840). Quantitative PCR was done using the iTaq Universal SYBR Green Supermix (Biorad, Cat: 1725121). All kits were used according to the manufacturer’s instructions. RNA was quantified using standard ddCT method and normalized to the endogenous control gene Hprt. A list of primer sequences used in this study are in Table S2.

### Statistical Analysis

All graphing of experimental data and statistical analyses were performed with GraphPad Prism (v11.0.0). Statistical tests performed for respective experiments are noted in the figure captions.

## Acknowledgements

We thank E. Ferrell and H. Dong for their help in colony maintenance and mouse breeding, and we thank O. Fedorova for synthesis and characterization of SLR14. We thank M. Linehan and P. Baevova for technical support, and P. Lu for discussion and guidance on the intranasal SLR14 stimulation. This work benefitted from data assembled by the ImmGen consortium.

## Author Contributions

S.A.E., A.C.H., S.B., A.I., A.M.P. designed the research; A.I., A.M.P., D.T.T., W.M.P., and R.A.F. designed and generated the RIG-I^mScar^ model; S.A.E., A.C.H., S.B., S.L., D.O.R., D.K., and T.M.M. performed the research; S.A.E., A.C.H., and S.B. performed data analysis; A.I. and A.M.P. supervised the research; S.A.E., A.C.H., S.B., A.I., and A.M.P. wrote the original manuscript; S.A.E., A.C.H., S.B., S.L., D.O.R., D.K., T.M.M., D.T.T., W.M.P., R.A.F., A.I., and A.M.P. provided manuscript revisions.

## Data Availability

All data used to inform the manuscript’s conclusions are present in the text and supplementary materials. Data will be made available upon reasonable request. The RIG-I^mScar^ mice are available from A.M.P. under a materials transfer agreement with HHMI.

---

**Supplementary Figure 1.**
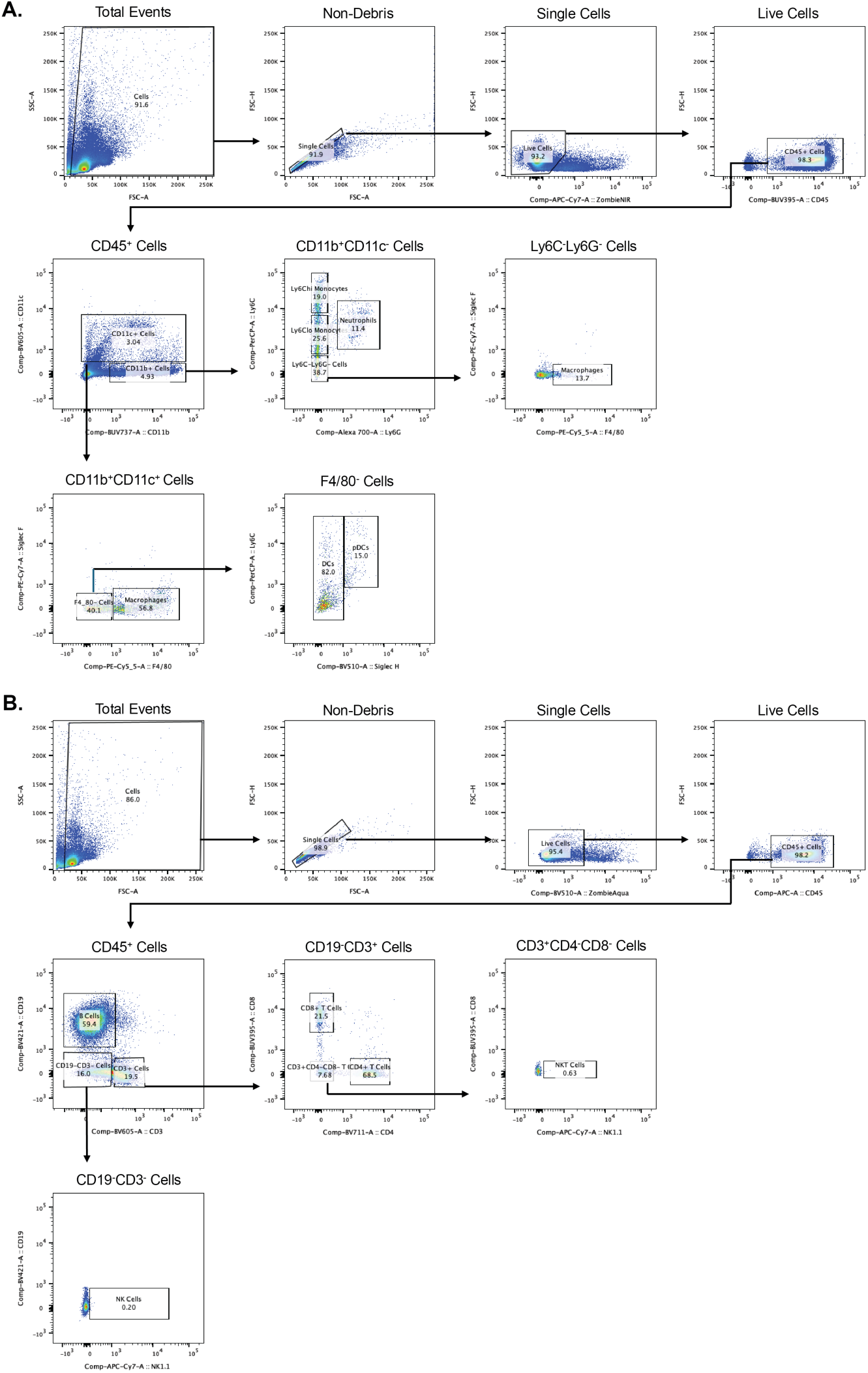
Flow cytometry gating strategies. Representative flow cytometry gating strategies of myeloid (A) and lymphoid (B) cells in the spleen.

**Supplementary Figure 2.**
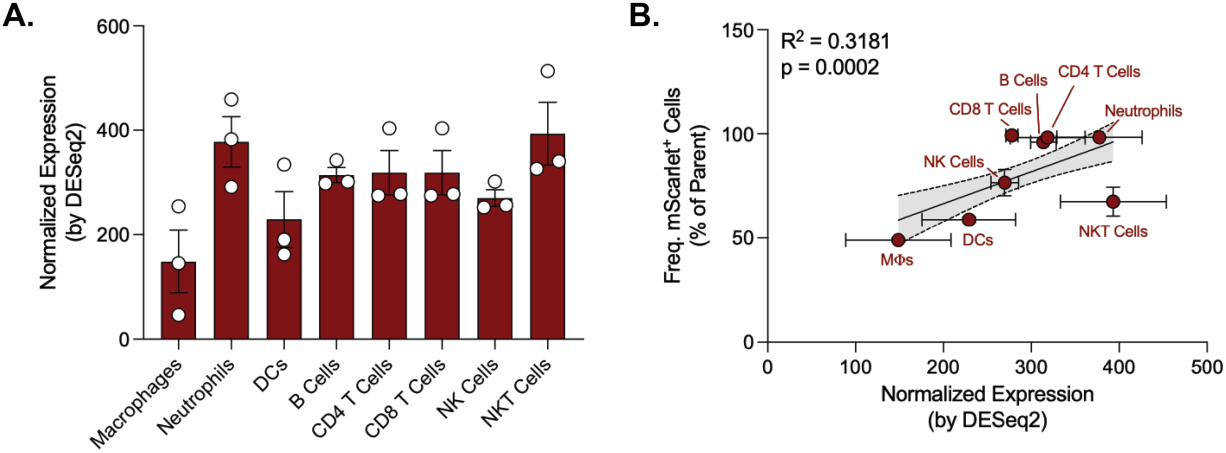
RIG-I^mScar^ signal correlates with single-cell transcriptomics. (A) Normalized transcript levels of RIG-I (Ddx58) in splenocytes from unstimulated, female C57BL/6 mice. Data were collected from the Immunological Genome Project (“ImmGen ULI: Male-Female Immune Differences”) and accessible in the GEO under GSE124829. Data are presented as mean ± SEM (n = 3). (B) Correlation of RIG-I^mScar^ signal in splenocytes with normalized transcript levels in the corresponding cell types, pulled from ImmGen as previously described. Correlation was calculated via simple linear regression.

**Supplementary Figure 3.**
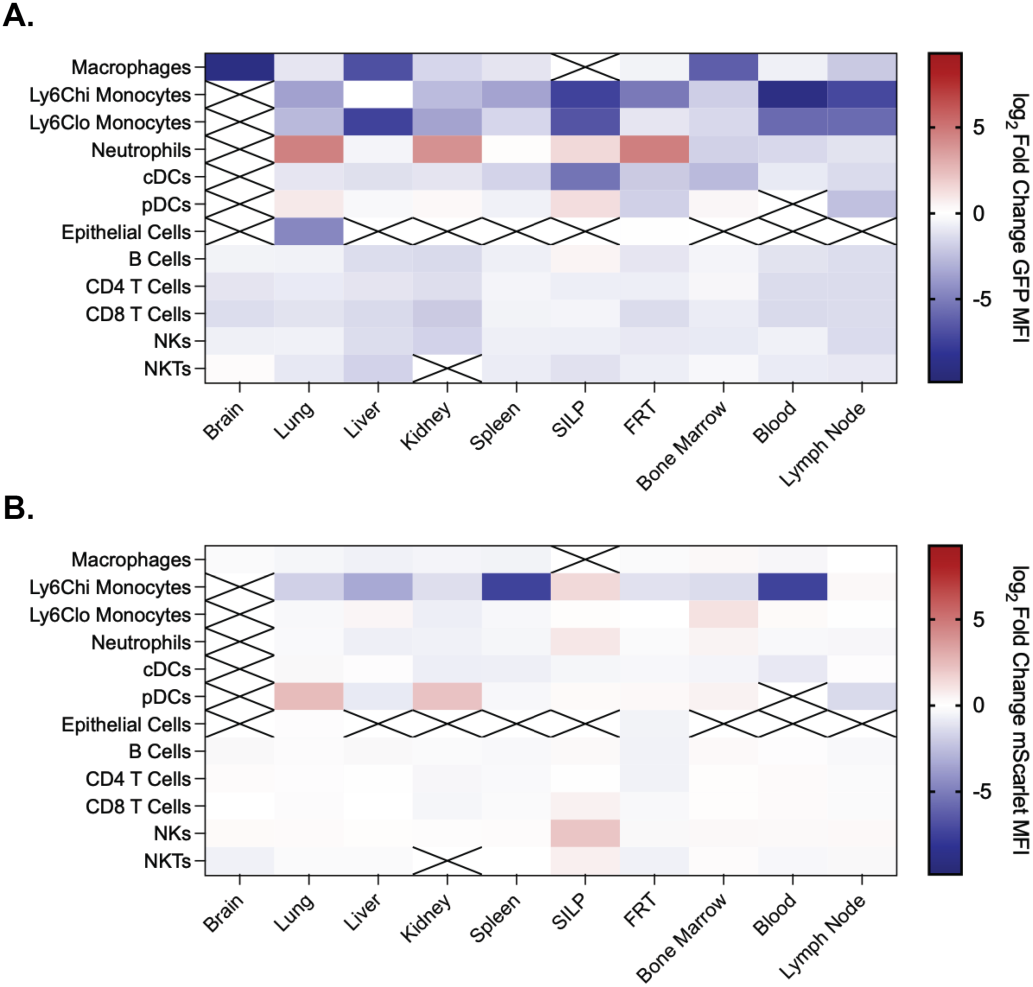
αIFNAR1 antibody treatment results in a global repression of Mx1^GFP^, but not RIG-I^mScar^, expression. Log_2_ foldchange in GFP (A) and mScarlet (B) MFI in the indicated cell types across tissues. Data are the same samples as in Fig. 2E-F. SILP = small intestine lamina propria; FRT = Female reproductive tract. cDCs = conventional dendritic cells; pDCs = plasmacytoid dendritic cells.

**Supplemental Figure 4.**
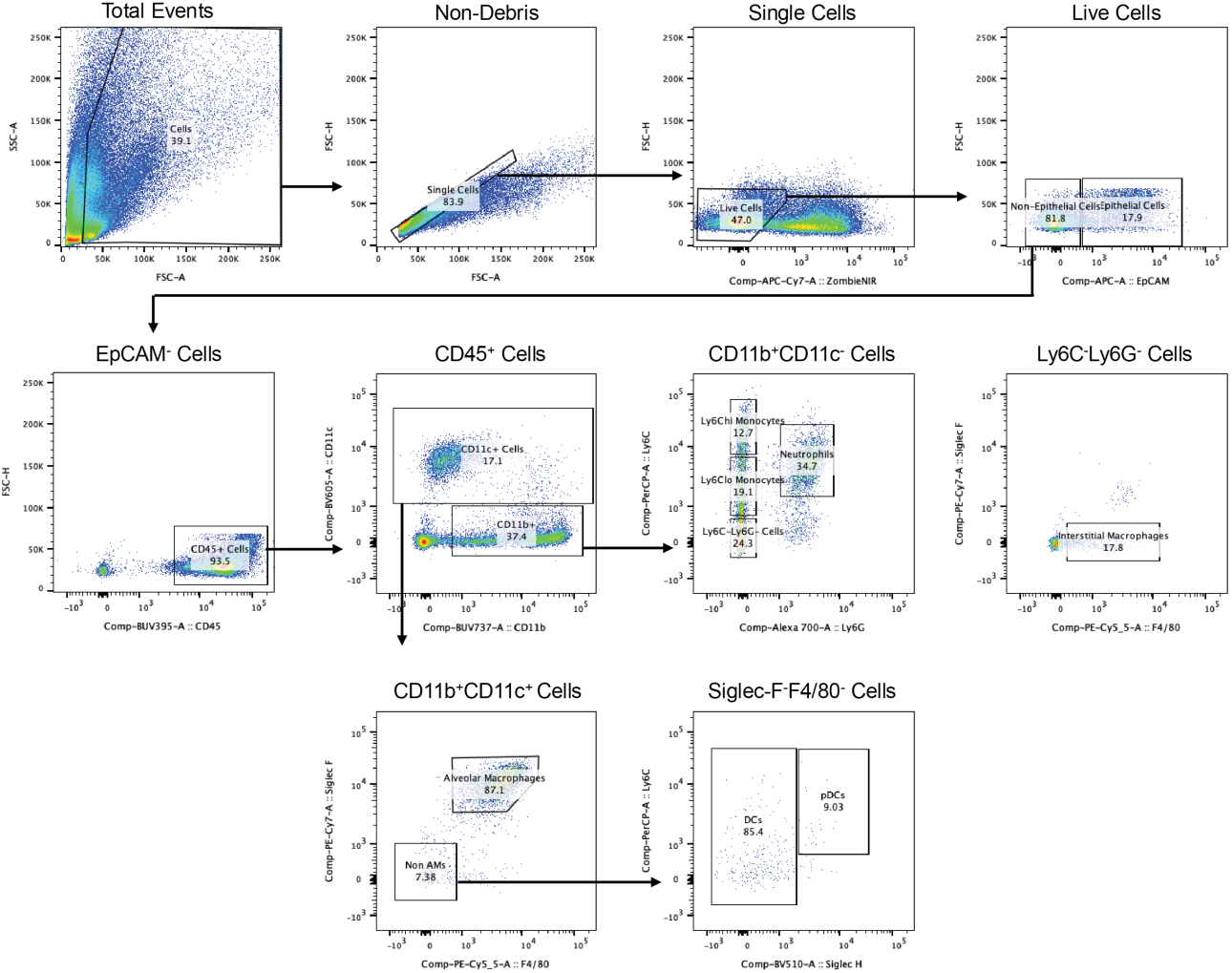
Representative flow gating strategy of lung cells.

**Supplementary Figure 5.**
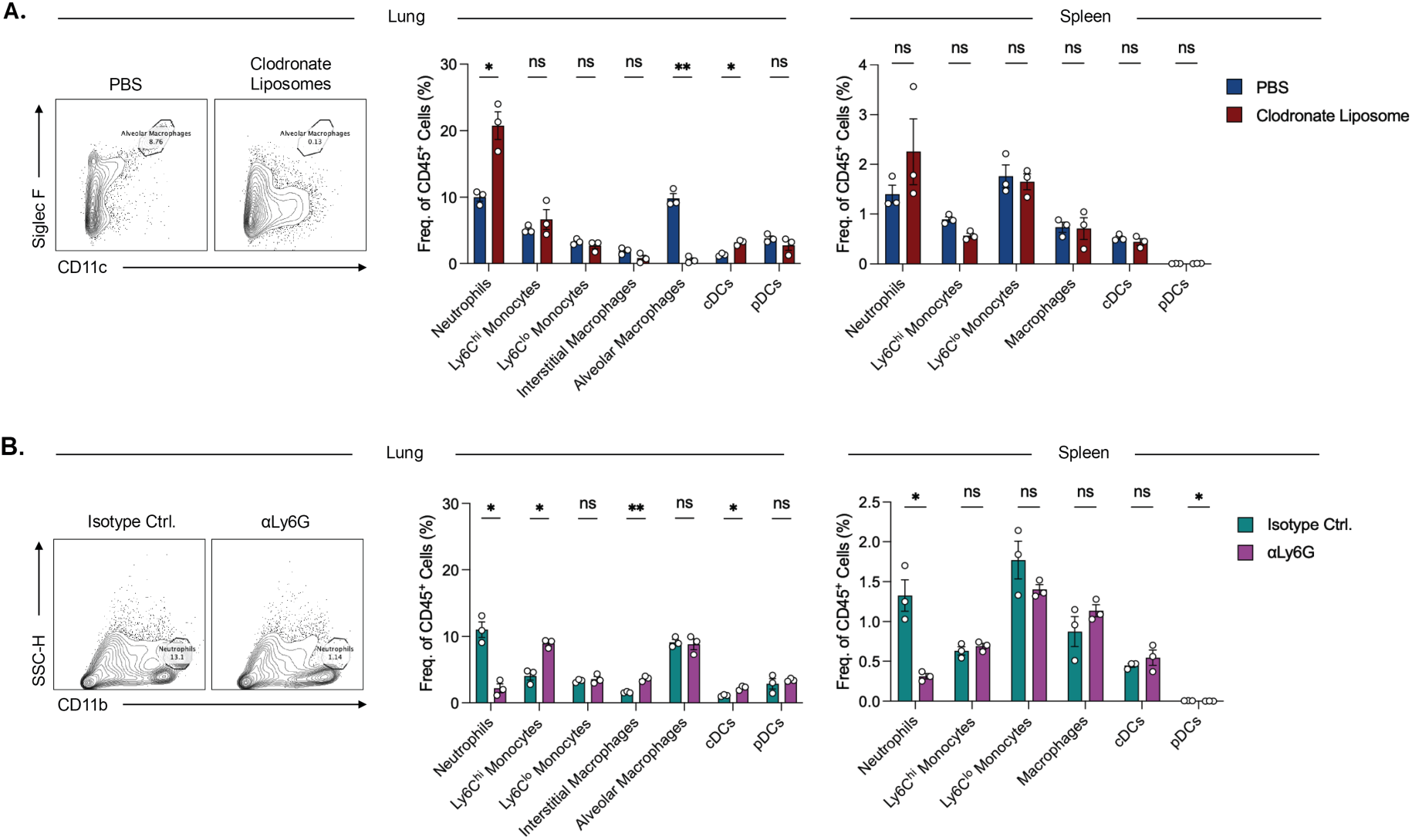
Successful depletion of lung neutrophils and alveolar macrophages by αLy6G antibodies and clodronate liposomes, respectively. (A) Representative flow cytometry plots of lung alveolar macrophages following PBS or clodronate liposome treatment as in Fig. 3B (left) and corresponding quantification of immune cell frequency in the lung (middle) and spleen (right). (B) Representative flow cytometry plots of lung neutrophils following isotype control or αLy6G-treatment as in Fig. 3E (left) and corresponding quantification of immune cell frequency in the lung (middle) and spleen (right). Data were analyzed via multiple unpaired t-tests with Holm-Šidák’s multiple comparisons test. All data are presented as mean ± SEM (n = 3/ea.). cDCs = conventional dendritic cells; pDCs = plasmacytoid dendritic cells.

**Supplemental Table 1.**
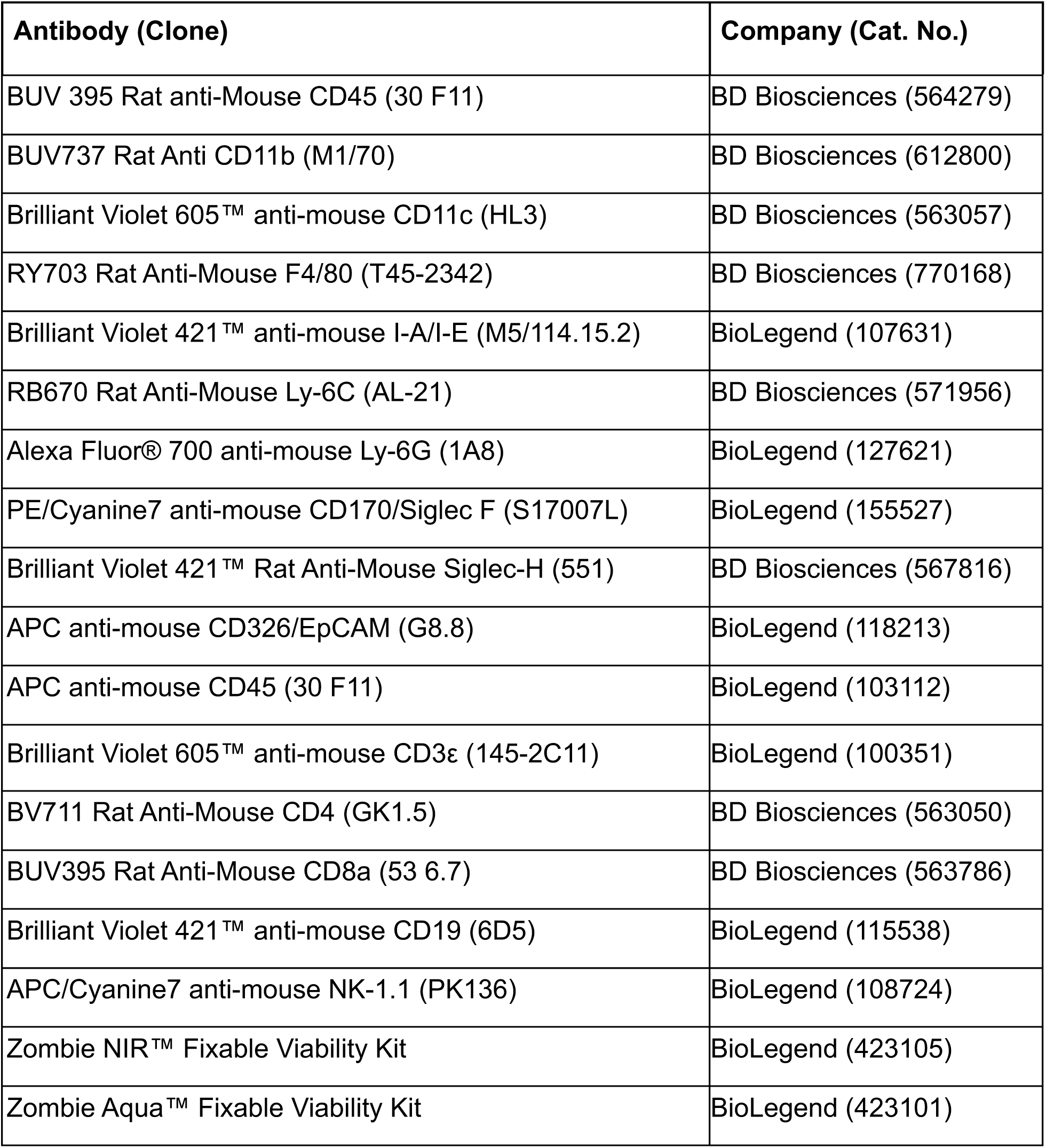
Flow antibodies used in the present study.

**Supplemental Table 2.**
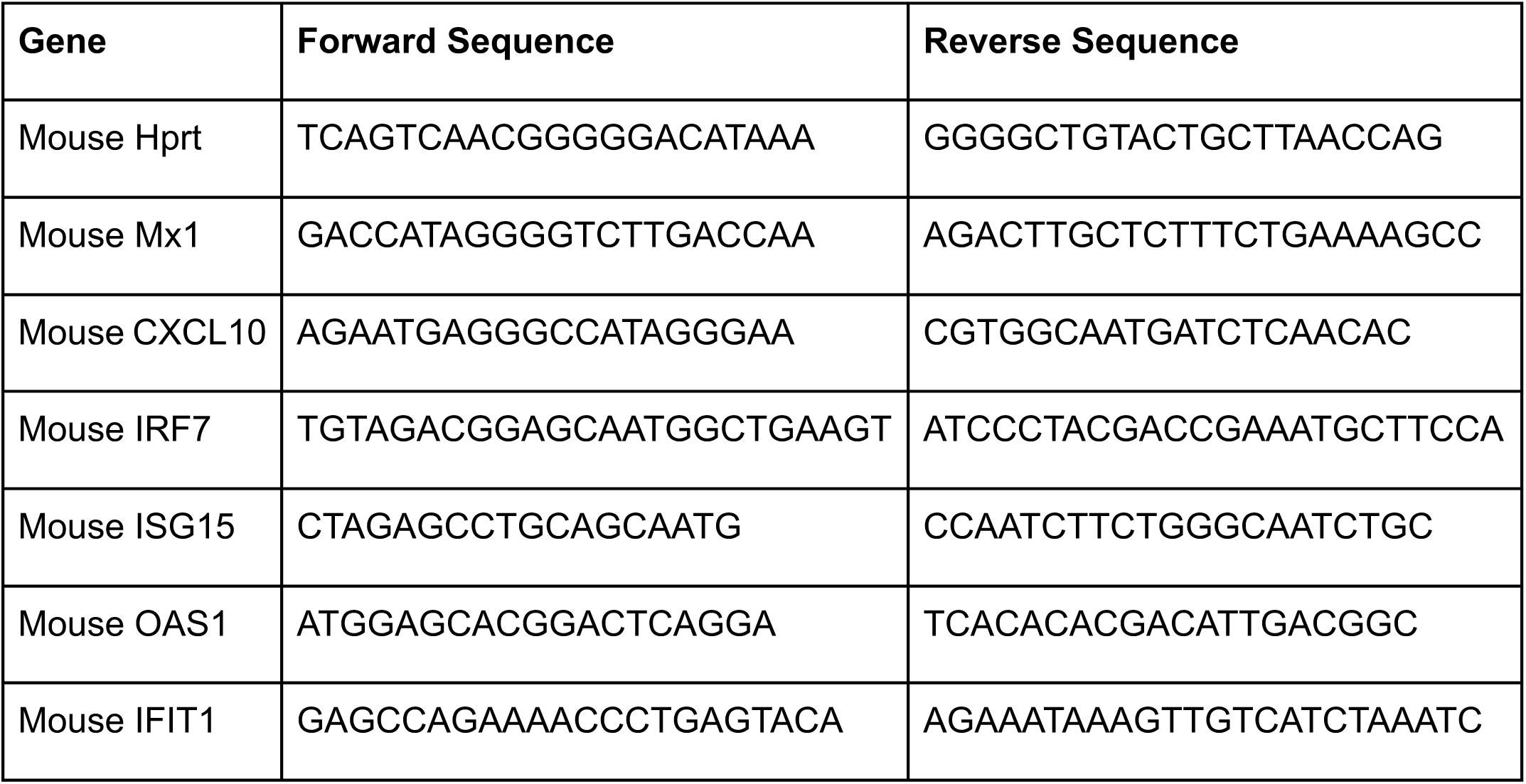
qPCR primers used in the present study.

